# Comparing the diversity and relative abundance of free and particle-associated aquatic viruses

**DOI:** 10.1101/2020.11.03.367664

**Authors:** Christine N. Palermo, Dylan W. Shea, Steven M. Short

**Affiliations:** Department of Physical and Environmental Sciences, University of Toronto Mississauga, Mississauga, Ontario, Canada; Department of Ecology and Evolutionary Biology, University of Toronto, Toronto, Ontario, Canada; Department of Biology, University of Toronto Mississauga, Mississauga, Ontario, Canada

**Author notes:** Address correspondence to Christine N. Palermo, or Steven M. Short,.

## Abstract

Metagenomics has enabled rapid increases in virus discovery, in turn permitting revisions of viral taxonomy and our understanding of the ecology of viruses and their hosts. Inspired by recent discoveries of large viruses prevalent in the environment, we re-assessed the longstanding approach of filtering water through small pore-size filters to separate viruses from cells before sequencing. We studied assembled contigs derived from < 0.45 μm and > 0.45 μm size fractions that were annotated as viral to determine the diversity and relative abundances of virus groups from each fraction. Virus communities were vastly different when comparing the size fractions, indicating that analysis of either fraction alone would provide only a partial perspective of environmental viruses. At the level of virus order/family we observed highly diverse and distinct virus communities in the > 0.45 μm size fractions, whereas the < 0.45 μm size fractions were comprised primarily of highly diverse Caudovirales. The relative abundances of Caudovirales for which hosts could be inferred varied widely between size fractions with higher relative abundances of cyanophages in the > 0.45 μm size fractions potentially indicating replication within cells during ongoing infections. Many of the *Mimiviridae* and *Phycodnaviridae*, and all *Iridoviridae* and *Poxviridae* were detected exclusively in the often disregarded > 0.45 μm size fractions. In addition to observing unique virus communities associated with each size fraction, we detected viruses common to both fractions and argue that these are candidates for further exploration because they may be the product of ongoing or recent lytic events.

**IMPORTANCE:** Most studies of aquatic virus communities analyze DNA sequences derived from the smaller, “free virus” size fraction. Our study demonstrates that analysis of virus communities using only the smaller size fraction can lead to erroneously low diversity estimates for many of the larger viruses such as *Mimiviridae, Phycodnaviridae, Iridoviridae*, and *Poxviridae*, whereas analyzing only the larger, > 0.45 μm size fraction can lead to underestimates of Caudovirales diversity and relative abundance. Similarly, our data shows that examining only the smaller size fraction can lead to underestimation of virophage and cyanophage relative abundances that could, in turn, cause researchers to assume their limited ecological importance. Given the considerable differences we observed in this study, we recommend cautious interpretations of environmental virus community assemblages and dynamics when based on metagenomic data derived from different size fractions.

## INTRODUCTION

Environmental viruses play key roles in shaping microbial communities and structuring ecosystems vital to the health of the planet and our species. They act as major drivers of microbial community succession (1, 2), influence global biogeochemical cycles (3), and may play important roles in the evolution of eukaryotes (4, 5). Although the Baltimore classification scheme (6) remains a valid system for categorizing fundamental groups of viruses, contemporary technologies have helped uncover a remarkable and diverse array of previously undiscovered viruses.

In particular, metagenomic data has rapidly expanded virus sequence databases and the documented diversity of viruses (7–15). Early metagenomic studies revealed the incredible diversity of viruses on a local scale, but limited global diversity, with the majority of viruses present at low abundances in the environment but acting as seed-banks for future infection (16). Additionally, metagenomics has allowed researchers to establish novel virus-host relationships on large scales with important ecological implications. For example, S. Roux et al. (12) identified > 12,000 virus-host relationships from publicly available metagenomes, assigning for the first time viruses to 13 bacterial phyla for which viruses had not been previously documented. Metagenomic time series data was also used to expand the range of *Mimiviridae* and ‘extended *Mimiviridae*’ hosts of virophages, and suggested that virophage hosts may include the *Phycodnaviridae* as well (17). K. Tominaga et al. (18) used *Bacteroidetes* metagenome-assembled genomes (MAGs) to augment existing reference databases and increase virus-host predictions by more than 3 times compared to using NCBI RefSeq alone.

Recently, metagenomic data has also become the primary source of novel virus genomes (19). Novel virophages were discovered in metagenomic datasets obtained from Organic Lake (20), Ace Lake (8), Yellowstone Lake (8, 21), Qinghai Lake (22), and Dishui Lake (23). The discovery rate of nucleocytoplasmic large DNA viruses (NCLDV) in particular has rapidly increased with data derived solely from metagenomic sequencing (14, 19, 24–26), generating several new taxonomic proposals up to the subfamily rank (27). As a result, the authority on virus classification, the International Committee on Taxonomy of Viruses (ICTV), has undergone a major pivot by accepting taxonomic proposals based on robust sequence data without the traditional requirements of cultured isolates, known host affiliations, or any other biological properties (28, 29). Clearly, metagenomics has fundamentally changed microbial ecology and continues to facilitate major discoveries, but as new analytical tools and pipelines are developed, constant exploration and scrutiny of new and existing bioinformatic and methodological approaches are necessary when addressing primary questions in environmental virology.

Early metagenomic studies of aquatic virus communities were conducted using DNA extracted from water samples passed through filters with 0.2 μm or smaller pore-sizes (30, 31). More recently, the discoveries of viruses with increasingly larger capsid sizes (as reviewed in J.-M. Claverie and C. Abergel (32)) necessitated re-evaluation of common filtration methods when attempting to capture entire aquatic virus communities. It is now clear that the < 0.2 μm size fraction excludes many diverse larger viruses, prompting some researchers to use 0.45 μm pore-size filtration in attempt to sample these larger viruses and obtain a more complete picture of the total virus community (33–36). For example, in one study of soils, use of 0.45 μm pore-size filters approximately doubled the number of observable virus-like particles compared to 0.2 μm pore-size filters (37). Thus, the choice of filtration schemes to enrich aquatic samples for viral community analysis is nontrivial, and is exacerbated by the presence of larger viruses, particularly those in the *Mimiviridae* family, that can be captured on 0.45 μm pore-size filters. Many of these larger *Mimiviridae* can be excluded from analyses of total virus communities when examining filtrates from 0.22 μm or 0.45 μm filtration, with the potential for slightly higher mimivirus recovery with 0.45 μm pore-size filters (37, 38). Additionally, viruses within or attached to cells or particles are likely to be captured on the filters and excluded from the analysis of filtrates used to obtain a picture of the viral community.

We previously studied virus communities in Hamilton Harbour (15), an urban eutrophic freshwater embayment of Lake Ontario, Canada that is well-studied due to its economic importance and long-term remediation efforts. We mined viral sequences from metagenome libraries derived from > 0.22 μm size fractions originally collected to study cellular communities. Typically, to avoid confounding viral and cellular sequences, viral sequences within this size fraction are filtered out and therefore, are not included in viral community analyses (39–42). Our study revealed diverse, seasonally dynamic viral communities that were distinct at different sites within the harbour. In most samples, virophages and *Mimiviridae* were highly abundant, contrasting other studies of similar environments that focused on viral communities in the < 0.22 μm size fraction. This has prompted us to examine both < 0.45 μm and > 0.45 μm size fractions collected from the same water samples in attempt to capture the entire virus community and to evaluate the diversity and relative abundances of different groups of viruses partitioning into each fraction.

## RESULTS

### Water filtration and DNA yields

Approximately 11 L of water from each site was filtered through a GC50 filter placed atop a PVDF filter. The entire volume of filtrate was incubated with FeCl_3_, but due to filters clogging, 2 PDVF filters per site were used to filter only 7 L of the FeCl_3_ filtrate. Therefore, approximately 4 L of water containing FeCl_3_ flocculates was discarded from each sample. DNA extractions from the < 0.45 μm size fractions yielded 4.9 – 8.0 μg of DNA at concentrations ranging from 61.2 – 99.6 ng/μl. For the > 0.45 μm size fractions, DNA was extracted separately from the PVDF and GC50 filters. DNA extracted from the PVDF filters yielded 0.5 – 0.7 μg of DNA at concentrations ranging from 5.4 – 6.5 ng/μl DNA, whereas DNA extracted from the GC50 filters yielded 3.0 – 4.0 μg of DNA per quarter of each filter at concentrations ranging from 30.2 – 39.5 ng/μl. After combining the extracted DNA from PVDF and GC50 filters, the final DNA concentrations for the > 0.45 μm size fractions ranged from 25 – 34 ng/μl; however, upon re-measurement at the sequencing centre, DNA concentrations used for sequencing library preparation were recorded in the range of 34.0 – 45.8 ng/μl (Table 1).

**Table 1:** Summary of sample parameters throughout the processing pipeline.

| **Sample** | **Initial DNA**<br>**concentration**<br>**(ng/μl)** | **Library**<br>**concentration**<br>**(ng/μl)** | **Reads**<br>**pre-QC** | **Reads**<br>**post-QC** | **Contigs**<br>**post-assembly**<br>**(mean length)** | **Contigs**<br>**assigned** |
| --- | --- | --- | --- | --- | --- | --- |
| < 0.45 μm<br>Site 1 | 61.2 | 7.22 | 34,682,792 | 26,909,942 | 350,572<br>(699) | 102,574 |
| < 0.45 μm<br>Site 2 | 82.7 | 6.53 | 33,438,900 | 25,831,282 | 329,713<br>(714) | 97,968 |
| < 0.45 μm<br>Site 3 | 99.6 | 6.07 | 36,189,024 | 28,542,680 | 377,348<br>(745) | 134,843 |
| > 0.45 μm<br>Site 1 | 43.8 | 4.24 | 35,222,034 | 25,937,694 | 290,644<br>(714) | 164,583 |
| > 0.45 μm<br>Site 2 | 34.0 | 5.82 | 37,718,578 | 28,401,490 | 356,847<br>(756) | 217,580 |
| > 0.45 μm<br>Site 3 | 45.8 | 5.27 | 40,863,560 | 28,521,306 | 192,849<br>(537) | 45,468 |

### Virus community composition in < 0.45 μm and > 0.45 μm size fractions

The majority of annotated contigs in the < 0.45 and > 0.45 μm size fractions were attributed to bacteria. Contigs annotated as viruses comprised 15% to 20% of all contig annotations in the < 0.45 μm size fractions, while fewer than 1 % of annotated contigs were of viral origin in the > 0.45 μm size fractions (Fig. 1). Diverse virus contigs were observed in both size fractions belonging to the virus groups Caudovirales, virophages (*Lavidaviridae*), *Mimiviridae*, *Phycodnaviridae*, other bacterial viruses, other dsDNA viruses, ssDNA viruses, and unclassified viruses. The < 0.45 μm size fractions were similar at the 3 sites and clustered closely together in the Bray-Curtis dissimilarity dendrogram (Fig. 1). While similar virus groups were found in both size fractions, most virus groups were present at only low relative abundances in the < 0.45 μm size fractions; Caudovirales, other bacterial viruses, and unclassified viruses comprised > 99% of all virus contigs in the smaller size fractions. Caudovirales relative abundances were remarkably consistent in the smaller size fractions at 82%, but were more variable in the larger size, fractions ranging from 2% to 42%. Additionally, there was more variability in the relative abundances of viral groups between sites in the > 0.45 μm size fractions than the < 0.45 μm size fractions (Fig. 1).

**Figure 1:**
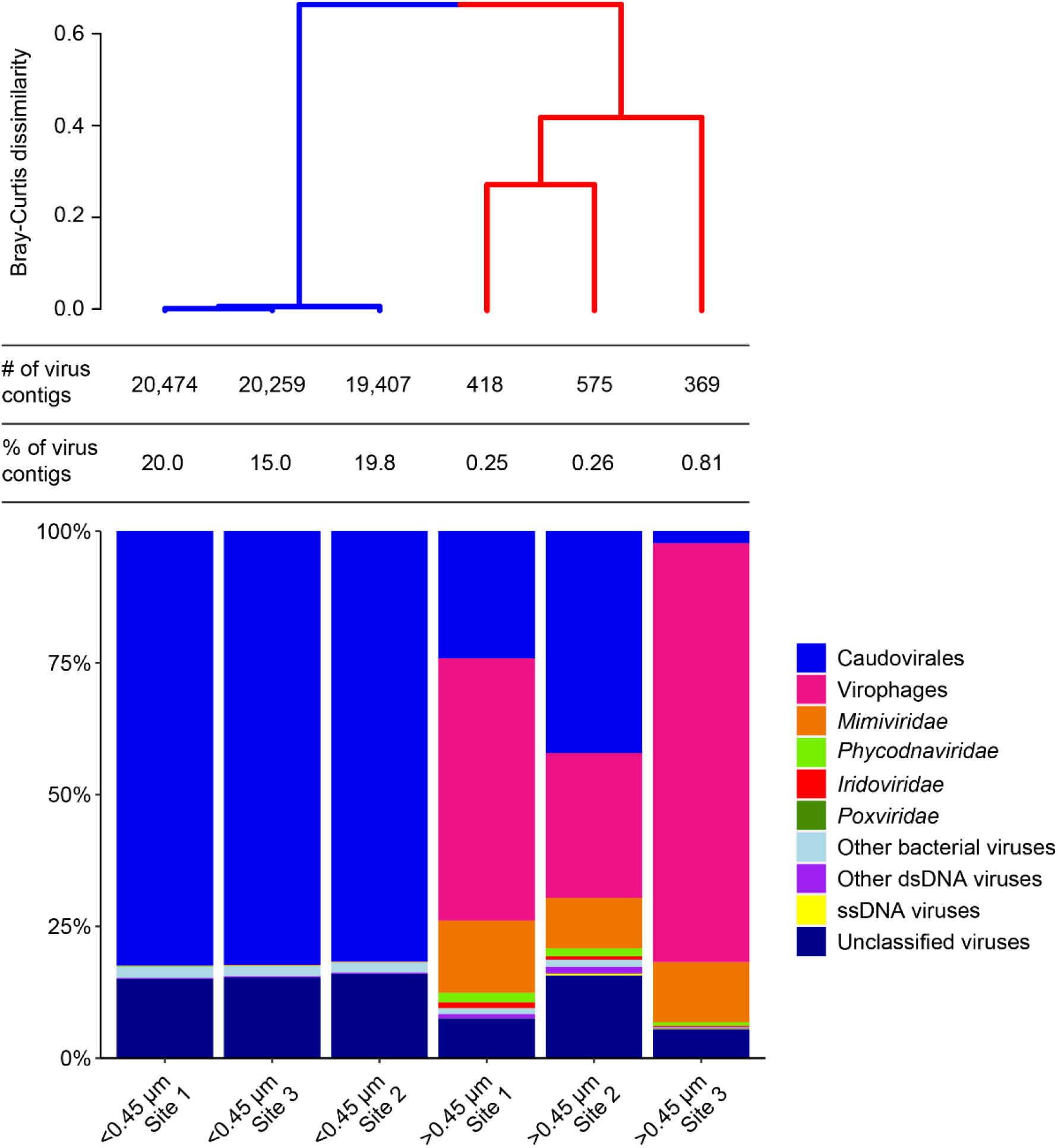
Relative abundances of virus groups from the < 0.45 μm size fractions and > 0.45 μm size fractions at each site. The table above the bars shows the number of virus contigs that comprise each bar and the percent of all contigs that were annotated as viral for each sample. Sample similarity is represented by the Bray-Curtis dissimilarity dendrogram above the table.

Relative abundances of virophages in the > 0.45 μm size fractions ranged from 27% to 79%, whereas *Mimiviridae* relative abundances ranged from 10% to 14%. In contrast, in the < 0.45 μm size fractions, virophage and *Mimiviridae* relative abundances were < 0.1% at all 3 sites. Algal viruses in the family *Phycodnaviridae* were present in all samples but at low relative abundances. In the < 0.45 μm size fractions, *Phycodnaviridae* were only 0.1% of the virus community at all 3 sites, but were slightly more abundant in the > 0.45 μm samples ranging from 1% to 2% relative abundance.

### Caudovirales community composition and host inference

*Siphoviridae* were the dominant Caudovirales family in all samples (Fig. 2). In the < 0.45 μm size fractions, the *Siphoviridae* comprised > 88% of all Caudovirales at the 3 sites, while the relative abundances of *Siphoviridae* in the > 0.45 μm size fractions ranged from 54% to 76%. *Myoviridae* were the second most abundant family in the > 0.45 μm size fractions, ranging in relative abundance from 14% to 36%, whereas the Podoviridae were the second most abundant family in the < 0.45 μm size fraction ranging from 4% to 5% relative abundance. Overall, *Podoviridae* were present at low relative abundances in every sample, but with slightly higher values in the < 0.45 μm size fractions. As shown in the Bray-Curtis dissimilarity dendrogram, the smaller size fractions were similar in community composition across sites whereas in the larger size fractions only Sites 1 and 2 clustered together; Site 3 was more similar to the smaller size fractions than to the other larger size fractions (Fig. 2).

**Figure 2:**
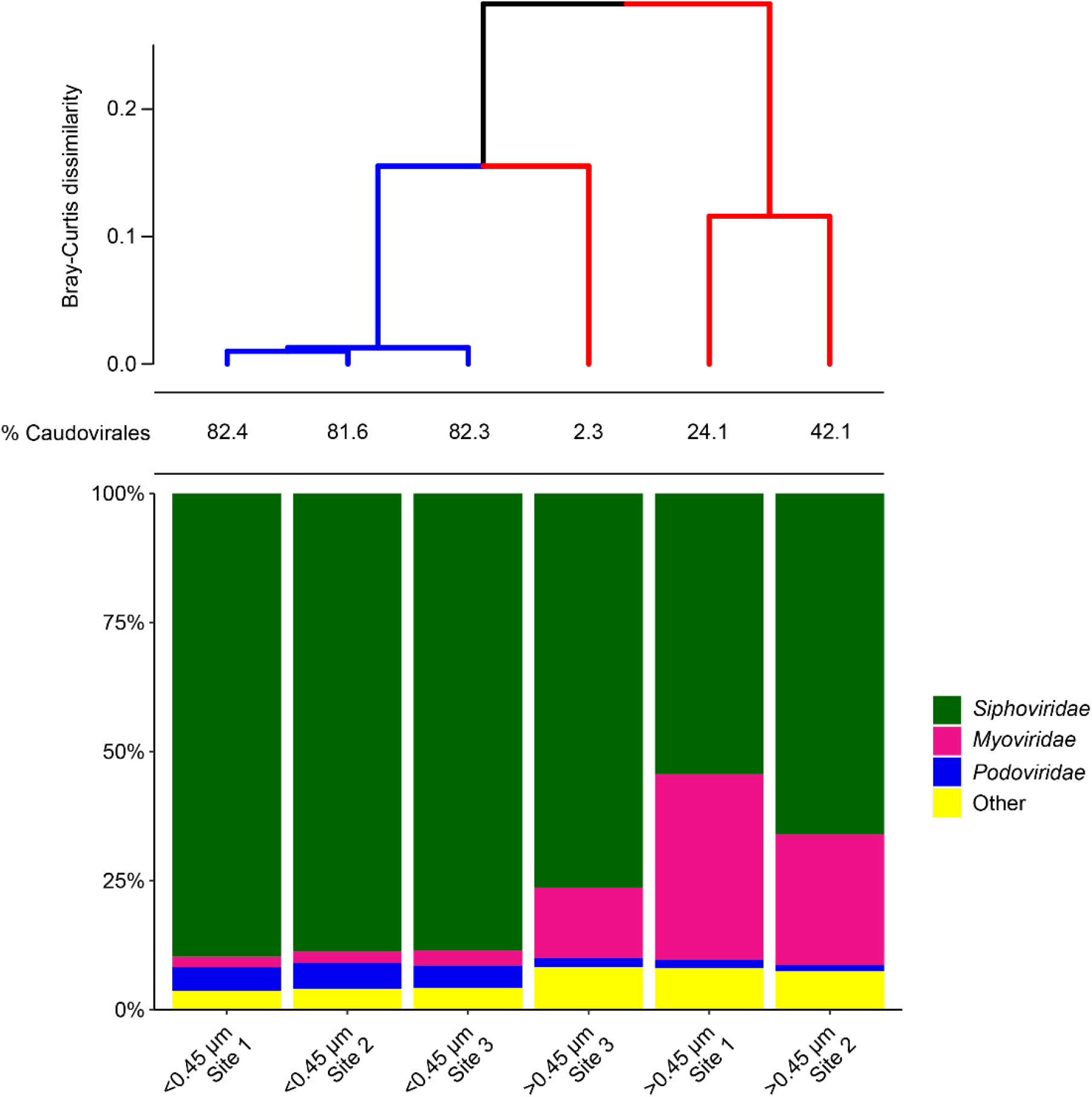
Relative abundances of Caudovirales families from the < 0.45 μm size fractions and > 0.45 μm size fractions at each site. Numerical values above the bars are the percent of Caudovirales in the entire virus community for each sample. Sample similarity is represented by the Bray-Curtis dissimilarity dendrogram.

Caudovirales with hosts that could be inferred from the virus name were sorted into cyanophages and non-cyanophages. Of the inferred cyanophages, phages of *Synechococcus* were the most abundant in the < 0.45 μm size fractions, ranging from 73% to 77% relative abundance (Fig. 3). In the > 0.45 μm size fraction at Site 3, *Synechococcus* was the most abundant inferred cyanophage host, whereas the > 0.45 μm size fractions at Sites 1 and 2 were dominated by phages of *Microcystis*. Phages of *Aphanizomenon* were detected in every sample, with higher relative abundances in the larger size fractions than the smaller size fractions. Like the clustering of Caudovirales families, all smaller size fractions clustered together, while the larger size fraction at Site 3 was more similar to the smaller size fractions than to the other larger size fractions (Fig. 3).

**Figure 3:**
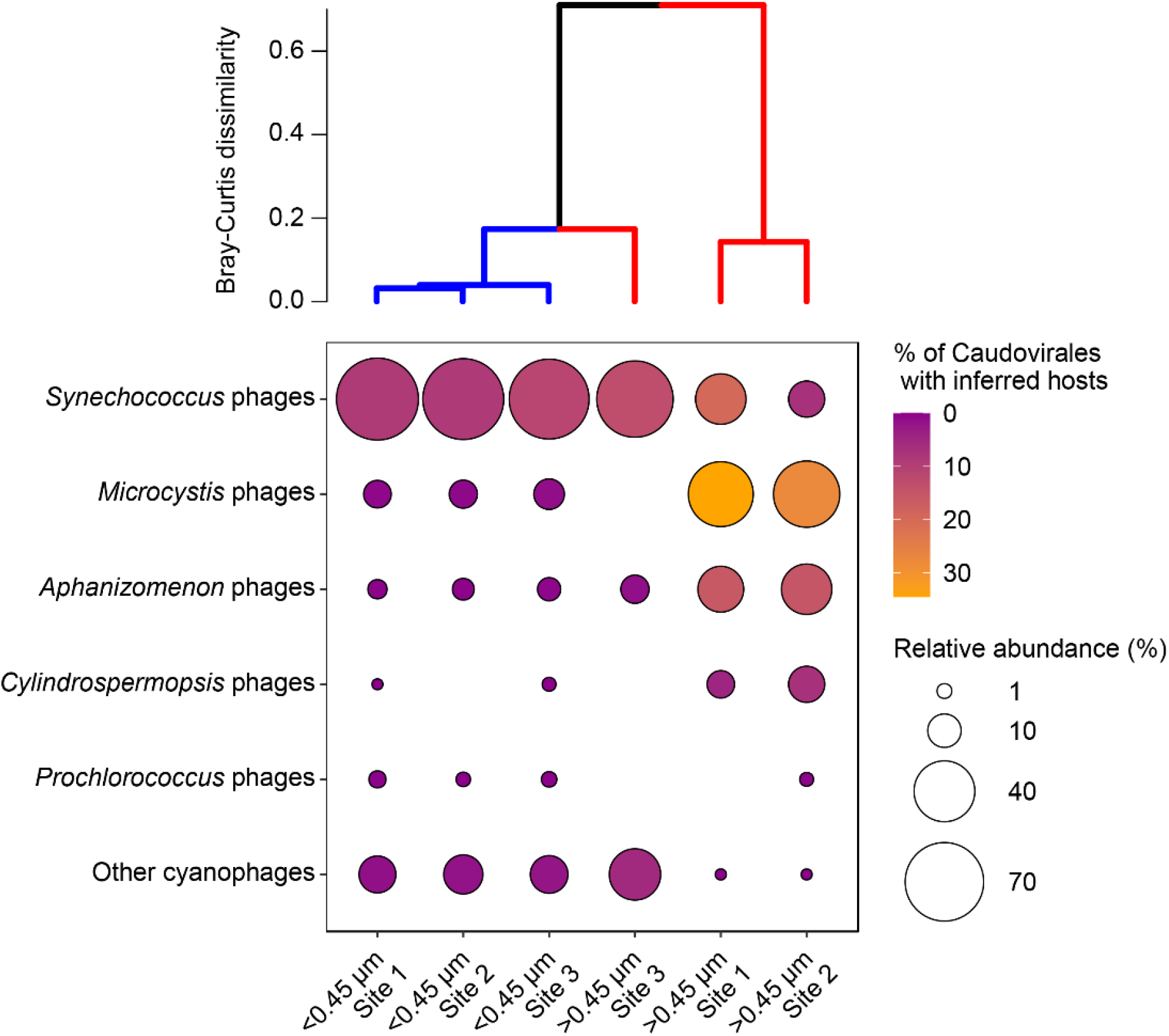
Bubble plot showing relative abundances of cyanophages for which hosts could be inferred with abundances denoted by the size of the bubble. Shading represents the percent of all Caudovirales with inferred hosts that each bubble represents. Sample similarity is represented by the Bray-Curtis dissimilarity dendrogram.

Among Caudovirales with inferred hosts other than cyanobacteria, phages of *Methylophilaceae*, Methylophilales and *Pontimonas* were most abundant in the < 0.45 μm size fractions, together comprising 53% to 60% of all Caudovirales with inferred hosts (Fig. 4). L5 viruses infect *Mycobacterium* and *Rhodococcus* species and were the most abundant phages of heterotrophic bacteria in the > 0.45 μm size fractions at Sites 1 and 2. Phages inferred to infect *Lactococcus* were the most abundant phages in the > 0.45 μm size fraction at Site 3, comprising 16% of all Caudovirales annotations in that sample. Across all samples, the “Other bacteriophages” category contained 85 unique phages, many of which were viruses of heterotrophic bacteria. In the Bray-Curtis dissimilarity analysis, distinct clusters formed for the < 0.45 μm size fractions and the > 0.45 μm size fractions. For both size fractions, Caudovirales communities from Sites 1 and 2 clustered together and the communities at Site 3 were less similar to either Site 1 or 2.

**Figure 4:**
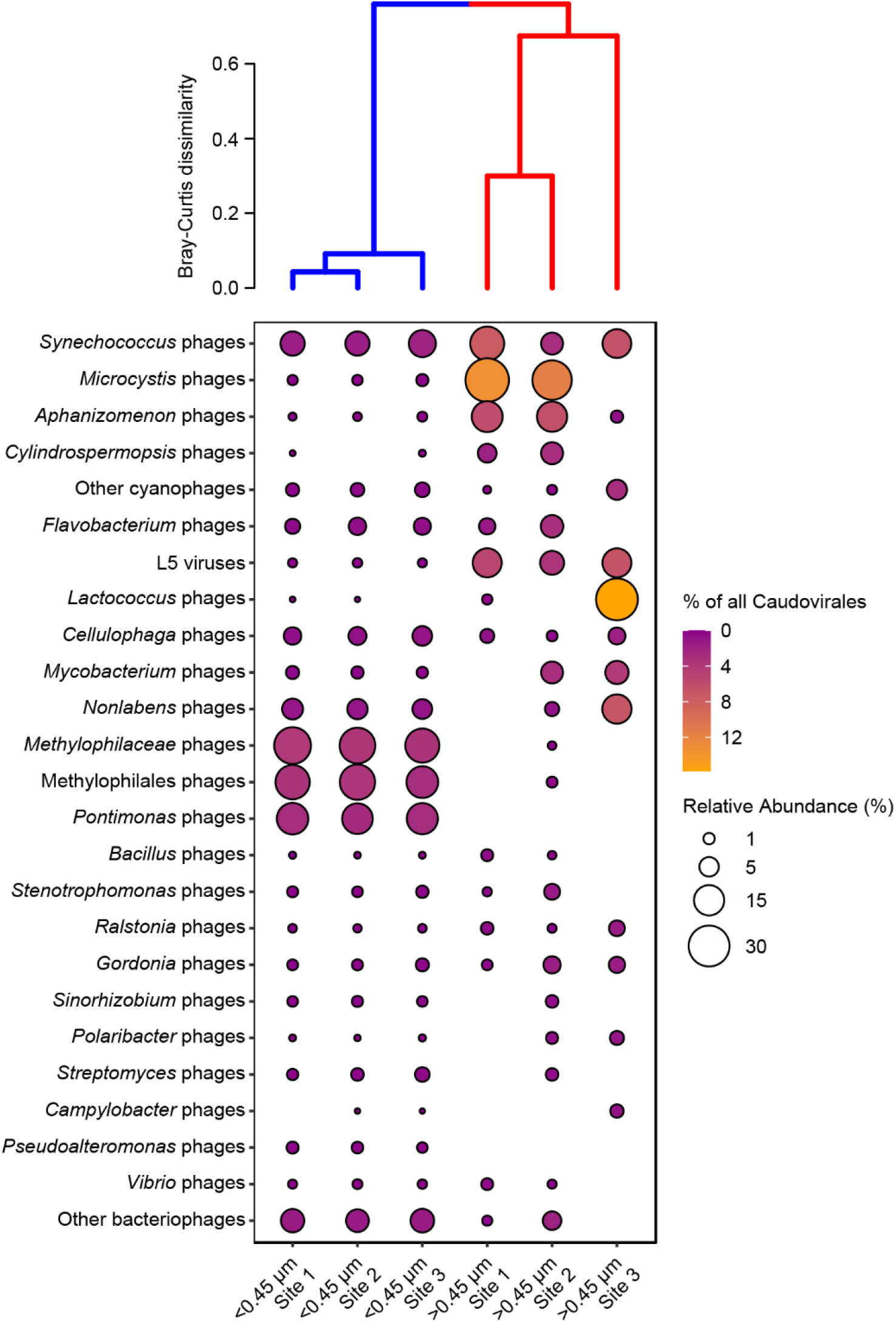
Bubble plot showing relative abundances of Caudovirales for which hosts could be inferred with abundances denoted by the size of the bubble. Shading represents the percent of all Caudovirales that each bubble represents. Sample similarity is represented by the Bray-Curtis dissimilarity dendrogram.

### Virus diversity in the < 0.45 μm and > 0.45 μm size fractions

Of 694 unique contig assignments for sequences generated from all 6 samples, 93% were detected in at least one sample from the < 0.45 μm size fractions, whereas only 24% were detected in at least one of the > 0.45 μm size fractions (Fig. 5a). 76% of unique contig assignments were found only in the < 0.45 μm size fractions, while 7% of unique contig assignments were found only in the > 0.45 μm size fractions. The two size fractions shared 17% of unique contig assignments, most of which were Caudovirales. 91 % of the unique contig assignments that were identified solely in the < 0.45 μm size fractions were Caudovirales, compared to 35% in the > 0.45 μm size fractions. The majority of unique Caudovirales and other bacterial viruses were detected only in the < 0.45 size fractions, whereas *Iridoviridae* and *Poxviridae* were only detected in the > 0.45 μm size fractions. In addition to the 11 unique *Mimiviridae* contigs detected in both size fractions, 7 unique *Mimiviridae* contigs were detected in the < 0.45 μm size fractions and 11 unique *Mimiviridae* contigs were detected in the > 0.45 μm size fractions. There were 18 unique *Phycodnaviridae* contigs detected only in the < 0.45 μm size fraction, compared to 6 unique *Phycodnaviridae* contigs detected only in the > 0.45 μm size fraction. Seven unique virophage contigs were detected in both size fractions and an additional 3 were detected in the < 0.45 μm size fractions.

**Figure 5:**
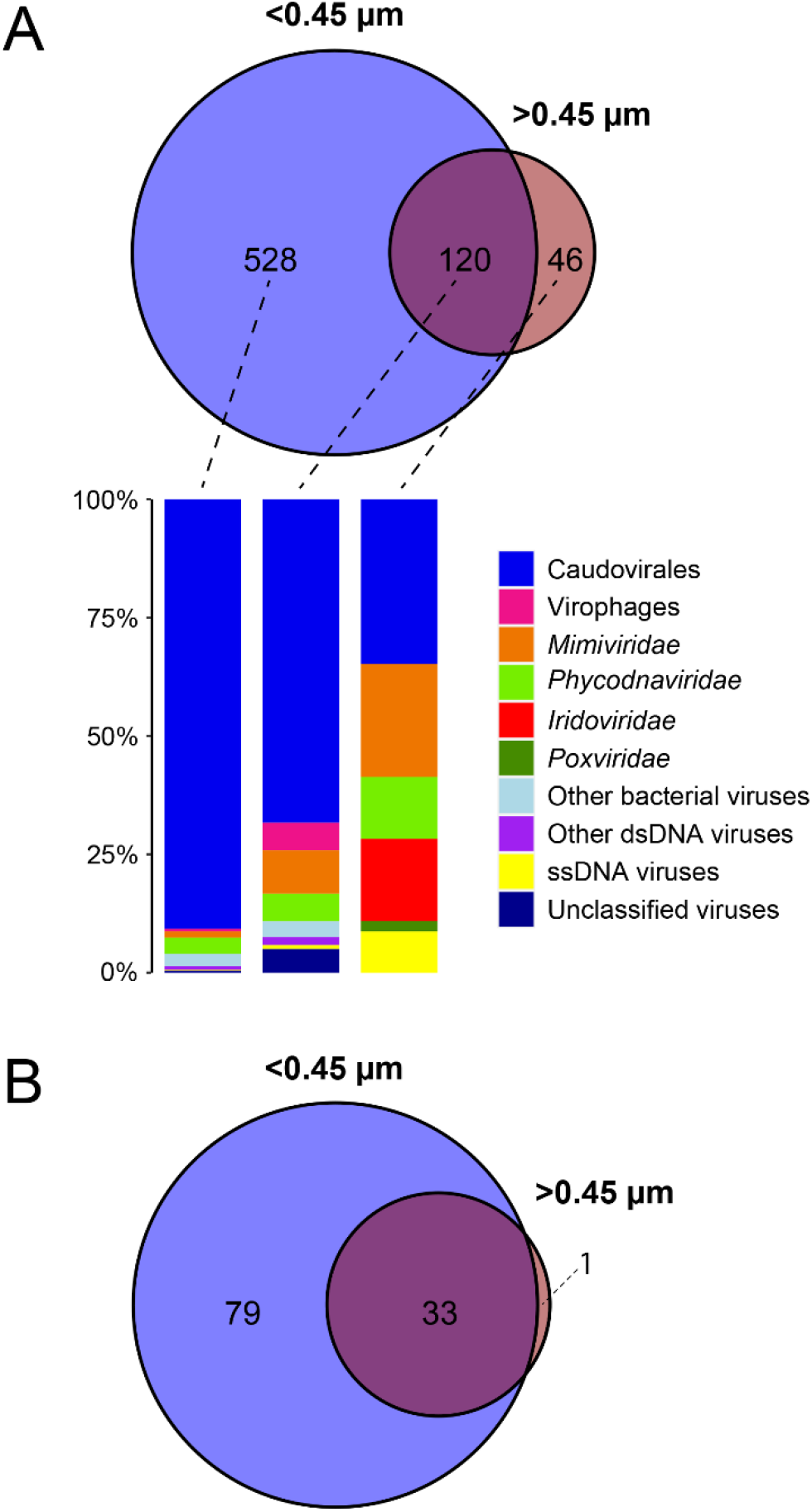
Unique and shared elements in the samples pooled into < 0.45 μm and > 0.45 μm size fractions. A) Venn diagram of the number of unique contig annotations in each size fraction. Bar charts show the relative abundance of virus groups in each category of the Venn diagram. B) Venn diagram of the number of unique inferred Caudovirales hosts in each size fraction.

There were 86, 87 and 89 unique Caudovirales with inferred hosts in the smaller size fractions at Sites 1, 2 and 3, respectively. We observed lower richness in the larger size fractions, which contained 15, 31 and 12 unique Caudovirales with inferred hosts at Sites 1, 2 and 3, respectively. All inferred Caudovirales hosts except one were captured in the < 0.45 μm size fractions (Fig. 5b). *Salinibacter* phage was the only Caudovirales with an inferred host that was found in one of the larger size fractions but not in any of the smaller size fractions. All Caudovirales with inferred hosts with > 1% relative abundance (i.e., those in Fig. 4) were captured by both size fractions, except *Pontimonas* and *Pseudoalteromonas* phages, which were not detected in the larger size fractions. Many of the 85 viruses with inferred hosts within the “Other bacteriophages” category were only detected in the smaller size fraction, and at low relative abundances.

Canonical correspondence analyses (CCA) were performed to assess the influence of physiochemical parameters on virus community composition. However, none of the tested parameters significantly explained the differences the data, including temperature, conductivity, dissolved O_2_, chlorophyll a, photosynthetically active radiation (PAR), salinity, pH, and Secchi depth. Rather, the different size fractions and host communities at each site were more important factors in explaining the variation in virus community diversity and relative abundances between samples.

## DISCUSSION

### Data considerations and assumptions

Beyond factors related to data processing programs and pipelines, there are considerations and assumptions related to our methods and results that merit some discussion. The sequencing methods used in this study only capture DNA viruses, and thus, the dsRNA and ssRNA virus communities were not considered. Furthermore, sequencing would only capture ssDNA viruses in dsDNA form, for instance during replication inside their hosts. This is reflected in our results, which showed that most ssDNA viruses were detected in the cell- and particle-associated > 0.45 μm size fractions. Therefore, the data and conclusions presented in this study apply almost exclusively to the dsDNA virus communities.

Compared to prokaryote and eukaryote sequences, virus sequences are underrepresented in reference databases influencing the reported diversity of viruses in metagenomes (discussed in detail in C. N. Palermo et al. (15)). This inevitably leads to a large portion of sequences in environmental metagenomes remaining unclassified and being discarded at the annotation step in the data processing pipeline. In order to increase the likelihood of annotating virus sequences, we used the NCBI-nr database which encompasses reference sequences from both curated and non-curated data sources. Of the virus sequences available in reference databases, some groups are more comprehensively represented than others. For example, Caudovirales genomes are two orders of magnitude more numerous than virophages genomes in NCBI RefSeq (data from https://www.ncbi.nlm.nih.gov/genomes/GenomesGroup.cgi). This leads to a higher likelihood of annotating Caudovirales sequences as originating from discrete taxa compared to virophage sequences. Our observations of disproportionally high virophage abundances compared to reference databases aligns with previous observations of their high relative abundances in larger, cell-associated size fractions (15, 17), and reinforces the notion that sequence annotations are only as good as the databases from which they are derived.

In this study we inferred the presence and abundance of viruses using the normalized relative abundances of assembled contigs. There are many viral genes present in prokaryotic and eukaryotic genomes due to the intimate evolutionary relationship between viruses and their hosts. For example, it is well known that many bacterial genomes are littered with cryptic prophage sequences (55), and endogenized retrovirus sequences are a common and surprisingly abundant feature of mammalian genomes (56). Unless viral homologues within host genomes are specifically annotated as such, they could be incorrectly annotated as viruses (57). Furthermore, metagenomic data cannot distinguish between viruses that are involved in active infections and inactive viruses that are adsorbed to cells or particles, which could also lead to an overestimation of viral influence. Therefore, a caveat of this study is the fact that we could not distinguish viral DNA integrated into host genomes versus viral DNA packaged within virions.

For the > 0.45 μm size fractions, DNA was extracted from 2 filter types: the 0.45 μm pore-size PVDF filters and the 0.5 μm nominal pore-size GC50 filters. The GC50 filters were placed on top of the 0.45 μm PVDF filters during water filtration, and therefore captured most of the biomass. This was evident from the thick layer of dark green biomass visible on the GC50 filters, whereas the PVDF filters below did not have any discolouration or visible biomass. For each site we combined the DNA from the two filter types proportionally based on the DNA concentrations from each of the filter extractions. For example, for Site 1, the DNA concentrations recovered from the 0.45 μm pore-size and GC50 filters were 6.5 ng/μl and 30.2 ng/μl corresponding to 18% and 82% of total DNA, respectively. Therefore, 7.2 μl of the extracted DNA from the 0.45 μm pore-size filter and 32.8 μl of the extracted DNA from the GC50 filter were combined. Rather than combining equal volumes or amounts of DNA from each filter, we chose to pool the DNA proportionally to avoid biasing the “total” DNA sample with DNA extracted from only one filter. Ultimately, the different amounts of DNA recovered from each filter type was proportionally represented in the DNA sample that was sent for sequencing.

Lastly, the methods of concentrating the viruses differed between the two size fractions, and therefore we were unable to distinguish the influence of size fraction versus virus concentration method on the diversity and relative abundance of the virus communities. For the > 0.45 μm size fractions, there was no additional concentration step beyond the filtration of water through the GC50 and PVDF filters. However, for the < 0.45 μm size fractions, comparable filter extraction methods are not possible. Filters with pore-sizes small enough to capture viruses have very low porosity allowing filtration of only small volumes and therefore, only minute amounts of DNA can be recovered from these filters, prohibiting effective library preparation for metagenomic sequencing. Therefore, for the < 0.45 μm size fractions, viruses were concentrated using FeCl_3_ flocculation (43). This approach was selected for its benefits over other widely used methods for concentrating aquatic viruses such tangential flow filtration (TFF), which is expensive, time consuming, and produces highly variable results (43). Furthermore, FeCl_3_ flocculation may be more comparable to filter-based extractions than TFF due to the capture and subsequent liberation of biomass from a solid surface. Although using a flocculation approach was an essential aspect of this study that allowed us to compare virus communities in the ‘particulate-associated’ (> 0.45 μm diameter) versus ‘free’ (< 0.45 μm diameter) fractions, it is possible that this method concentrated certain types of viruses more effectively than others, potentially influencing our estimates of virus diversity and relative abundance, although taxon-specific biases have not previously been identified with this viral recovery method.

### Comparisons of the < 0.45 μm and > 0.45 μm size fractions

We observed major differences between the < 0.45 μm size fractions and > 0.45 μm size fractions at almost all levels of analysis. DNA extractions from the smaller size fractions yielded more DNA than the larger size fractions. DNA was extracted from only a quarter of the filters for the larger size fractions, and DNA yield may have been impacted by incomplete lysis, adsorption to particles, or incomplete recovery of supernatant after bead beating. Sequencing library concentrations were higher in the smaller size fractions (Table 1), but despite this, there were no apparent trends in the number of reads generated before or after quality control, nor the total number of contigs assigned. The number of virus contigs assigned was much higher in the < 0.45 μm size fractions than the > 0.45 μm size fractions (Fig. 1), as expected due to the enrichment of virus particles in the < 0.45 μm size fractions.

For the sake of brevity, we will use only the virus name or group when referring to contigs annotated as those viruses. Across all levels of taxonomic analysis, the smaller size fractions clustered closely together in the Bray-Curtis dissimilarity dendrogram indicating few differences in the relative abundances of virus groups between sites. The larger size fractions had higher Bray-Curtis dissimilarity values at all levels of analysis. At the levels of Caudovirales families and Caudovirales with inferred cyanobacterial hosts, the larger size fraction from Site 3 clustered more closely with the smaller size fractions than with the other larger size fractions. At the order and family level, the smaller size fractions consisted almost entirely of Caudovirales, other bacterial viruses and unclassified viruses (Fig. 1) whereas the larger size fractions were more diverse with lower relative abundances of Caudovirales, but higher relative abundances of virophages, *Mimiviridae, Phycodnaviridae, Iridoviridae, Poxviridae*, other dsDNA viruses, and ssDNA viruses. Its also notable that the larger size fractions had wider ranging relative abundances of Caudovirales families (Fig. 2). While *Siphoviridae* were the most abundant Caudovirales family in every sample, relative abundances of *Myoviridae* in the larger size fractions were higher and more variable between the sites. Previous metagenomic studies of Lake Ontario and nearby Lake Erie found that *Myoviridae* were the dominant Caudovirales family (58). Likewise, metagenomic data from Lake Baikal showed a predominance of *Myoviridae* (59, 60), whereas *Siphoviridae* and *Myoviridae* relative abundances were similar in Lakes Michigan, Pavin, and Bourget (61, 62). Therefore, our findings of highly abundant *Siphoviridae* relative to *Myoviridae* in Hamilton Harbour contrasts previous studies of Caudovirales communities in other freshwater lakes.

Of the Caudovirales with inferred hosts, the larger size fractions from Sites 1 and 2 had higher relative abundances of cyanophages than the smaller size fractions and the Site 3 larger size fraction (Fig. 3). These higher relative abundances in the cell- and particle-associated size fractions may indicate active cyanophage infections, as Hamilton Harbour is known to experience algal blooms during the summer and late autumn (63).

*Microcystis* phage sequences were particularly high in relative abundance in the > 0.45 μm size fractions from Sites 1 and 2, suggesting ongoing active infections of *Microcystis* spp. *Microcystis* is a well-known bloom-forming species in Hamilton Harbour (64) and Lake Ontario (65), and therefore high relative abundances of *Microcystis* phages were expected. However, in the < 0.45 μm size fractions, contribution of *Microcystis* phages to the total cyanophage community was low, demonstrating the different insights into the virus communities that the two size fractions provide. Moreover, *Microcystis* phages were not detected in the > 0.45 μm size fraction from Site 3, reinforcing previous findings of distinct virus communities over relatively small spatial scales in Hamilton Harbour (15). We observed a more pronounced case of differences between the size fractions at Site 3, where *Lactococcus* phage sequences were highly abundant in the > 0.45 μm size fraction, but undetectable in the < 0.45 μm size fraction. Across all samples and Caudovirales with known hosts, the *Lactococcus* phages in the larger size fraction from Site 3 had the highest percent abundance of all Caudovirales annotated in that sample. The most abundant *Lactococcus* phage in this sample was *Lactococcus* phage 16802. This phage infects *Lactococcus lactis*, which was found to be the most abundant lactic acid bacterium year round in 7 Japanese lakes (66). The discovery of highly abundant *Lactococcus* phages in a eutrophic embayment of the Great Lakes is intriguing given that *Lactococcus* species may control harmful algal blooms by acting as cyanobacteriacidal bio-agents (67). Regardless of their potential ecological interactions with bloom forming species, the high relative abundance of *Lactococcus* phages is indicative of their ecological importance in this system.

In all the < 0.45 μm size fractions, phages infecting Methylophilales, *Methylophilaceae*, and *Pontimonas* were the most abundant Caudovirales with inferred hosts (Fig. 4). *Methylophilaceae* is a ubiquitous and metabolically diverse bacterial family (68) with observed enrichments in the Great Lakes surface waters (69). They may have major roles in global carbon cycling (70) and nitrogen cycling via methanol oxidation linked to denitrification (71), and seem to be ecologically important in Hamilton Harbour, a high nutrient and high biomass environment. *Methylophilaceae* grow rapidly and have high biomass yields (70), potentially leading to our observations of high relative abundances of their phages during a major lytic event.

The *Pontimonas* genus contains only one type strain, *Pontimonas salinivibrio* strain CL-TW6^T^, which was isolated from a coastal marine environment along with its virus. Its genome is streamlined with an exception of several toxin/antitoxin systems, and it was thus hypothesized to occupy environments that experience significant stresses (72), such as Hamilton Harbour. The researchers concluded that its lytic virus was poorly adapted to replicate in this strain since repeated attempts to sequence its genome were unsuccessful. Given the high abundances of this *Pontimonas* phage in all < 0.45 μm size fractions from Hamilton Harbour, it presumably replicates successfully within its host, yet its absence in all of the > 0.45 μm size fractions warrants further investigation of this virus-host system that may be prevalent in Hamilton Harbor.

Less than 3% of unique Caudovirales annotations were not detected in the < 0.45 μm size fractions (Fig. 5A), and only 1 Caudovirales with an inferred host was not observed in this fraction (Fig. 5B). These observations may indicate that in addition to capturing the most abundant Caudovirales with inferred hosts, the < 0.45 μm size fractions captured diverse Caudovirales, with inferred hosts, that were present at low relative abundances that may be missed in the > 0.45 μm size fractions. However, many of the other virus families, including *Phycodnaviridae, Mimiviridae, Iridoviridae*, and *Poxviridae*, were absent or poorly represented in the < 0.45 μm size fractions. Likely due to their large capsid sizes, the majority of *Mimiviridae* were found only in the > 0.45 μm size fractions (Fig. 5A). *Iridoviridae* and *Poxviridae* were only found in the > 0.45 μm size fractions despite having small enough capsid sizes to pass through the 0.45 μm filters if present in the water column as free viruses. ssDNA viruses were detected primarily in > 0.45 μm size fraction, likely in dsDNA form while replicating within cells. Therefore, the smaller size fractions effectively captured Caudovirales, whereas the larger size fractions were more effective at capturing the diversity of larger viruses in the samples.

Interestingly, a subset of viruses was detected in both size fractions (Fig. 5), suggesting their high activity and ecological importance in this environment. These viruses were sufficiently abundant to be detected as free viruses in the water column as well as in the particulate- and cell-associated fraction, potentially indicating ongoing or recent lytic events. Of particular importance to Hamilton Harbour, we detected a variety of cyanophages in both size fractions, including phages of Microcystis (Ma-LMM01, MaMV-DC), Prochlorococcus (P-HM2), Aphanizomenon (vB_AphaS-CL131), and Synechococcus (ACG-2014f, S-PM2, S-CAM7, S-T4, S-CBS2, S-SKS1, S-PRM1, S-CAM22, S-CBS4, S-WAM2, S-CAM8, S-TIM5, KBS-S-2A). We also detected viruses of eukaryotic primary producers in both size fractions, including *Chrysochromulina ericina* virus, unclassified Chloroviruses, Yellowstone Lake Phycodnavirus 2, and *Micromonas* sp. RCC1109 virus MpV1. Identifying overlapping virus communities in different size fractions provides the impetus for further exploration because these viruses may be the product of recent or ongoing, active viral infections in aquatic environments. The hypothesis that viruses found in both size fractions are the product of ongoing infection warrants further experimental investigations that were beyond the scope of this study, for example via concurrent DNA and RNA analyses to evaluate the activity of viruses detected in both size fractions.

### Site comparisons and similarities with previous study of Hamilton Harbour viruses

Across all levels of analysis and both size fractions, Sites 1 and 2 were generally more alike than Site 3. The most noticeable differences between sites were in sequences from the > 0.45 μm size fractions. For example, relative abundances of Caudovirales sequences in all < 0.45 μm size fractions were 82%, whereas Caudovirales relative abundances in the > 0.45 μm size fractions were 24%, 42%, and 2% at Sites 1, 2 and 3, respectively (Fig. 1). Similarly, virophage relative abundances were 0.1% in all < 0.45 μm size fractions, whereas virophage relative abundances in the > 0.45 μm size fractions were 50%, 27%, and 79% at Sites 1, 2 and 3, respectively. In the > 0.45 μm size fractions, Site 3 community profiles also differed from Sites 1 and 2 at the level of Caudovirales with inferred hosts. The most abundant cyanophage host at Site 3 was *Synechococcus*, which was also the case in all < 0.45 μm size fractions (Fig. 3). While Sites 1 and 2 were dominated by phages of *Microcystis*, Site 3 contained a high relative abundance of *Lactococcus* phages (Fig. 4).

As previously documented (15), we observed distinct virus communities over small spatial scales in Hamilton Harbour. Modelling of summer circulation patterns revealed the occurrence of 3 eddies in the harbour (73). Sites 1 and 2 are situated in the middle and at the edge, respectively, of the largest eddy in the middle of the harbour, whereas Site 3 is located within a smaller eddy in the western part of the harbour. The location of the sites with respect to these eddies is possible explanation for the different communities we observed, although water circulation patterns at the time of sampling were not assessed. Site 3 is also closer in proximity to wastewater inputs entering the harbour through Spencer Creek, which may stimulate microbial growth at this site, in turn shaping the virus communities. Whatever the driving factor, it is interesting that distinct virus communities were again observed over small spatial scales in Hamilton Harbour which reinforces our previous observations that this pattern of community structure is a stable ecological phenomenon rather than a chance, singular observation from a snap-shot study of the environment.

Additionally, there were other similarities between the virus communities observed in the > 0.45 μm size fractions in this study and the > 0.22 μm size fractions from Hamilton Harbour in 2015 (15). In 2015, Sites 1 and 3 were sampled biweekly from July to September. Different methodology was used to study cell- and particle-associated viruses in Hamilton Harbour in 2015 and 2019; there were major differences in water sampling, filtration, and DNA extraction, and the samples were collected and processed by different researchers several years apart. Furthermore, we used different sequencing machines, raw read lengths, and data analysis parameters. Despite these large differences in methodology, in both studies we observed similar patterns of high virophage abundances in the > 0.45 μm size fractions, which were particularly pronounced at Site 3. As in the 2015 samples, *Mimiviridae* relative abundances in the > 0.45 μm size fractions did not fluctuate to the same extent as the virophages, despite their presumably intimate association. Like the study of 2015 samples, in this study of samples from 2019 we observed low relative abundances of *Phycodnaviridae* even though Hamilton Harbour is known to support high algal biomass during the sampling period. Again, this indicates that *Mimiviridae* may be the dominant algae-infecting viruses in this system. As a final comment on the similarities of this study and our previous study (15), all physiochemical factors measured were poor explanatory variables of viral community composition and relative abundance, likely reflecting the diverse host ranges of most virus groups.

### Experimental and ecological relevance

In this study we opted to use 0.45 μm pore-size filters rather than 0.2 μm pore-size filters to maximize the likelihood of capturing large, free virus particles in the filtrate in attempt to distinguish between “free viruses” in the < 0.45 μm size fraction and “cell- or particle-associated viruses” in the > 0.45 μm size fraction. *Mimiviridae* and other large DNA viruses are an exception to this generalization, as they could be captured on the > 0.45 μm pore-size filters if present as free viruses in the water column. Traditional viromics studies that look only at the < 0.2 μm or < 0.45 μm size fractions may exclude large portions of the *Mimiviridae*, *Phycodnaviridae*, *Iridoviridae*, and *Poxviridae* communities, potentially leading to an underestimation of their influence on the microbial communities. Additionally, members of the Caudovirales community involved in ongoing infections may be overlooked. Conversely, mining viral sequences in the cell- and particle-associated size fractions may not sufficiently capture the diversity of Caudovirales, many of which may be present at low abundances but acting as a seed bank (16) for future infection. The > 0.45 μm size fractions captured the diversity of larger viruses more effectively than the < 0.45 μm size fractions; however, many viruses in the > 0.45 μm size fractions are likely missed by sequencing efforts due to the low representation of viruses in these datasets. Larger cellular genomes comprise most of the DNA in the > 0.45 μm size fractions, leading to low sequencing coverage of viruses, and therefore likely reducing diversity estimates by only permitting sequencing of the most abundant viruses. While all discrete virophage taxa were observed in the < 0.45 μm size fractions, virophage contigs observed in the > 0.45 μm size fractions were often longer and more numerous. Therefore, high virophage relative abundance estimates from the larger size fractions were not simply a result of low detection of other groups of viruses. The higher virophage relative abundances in the larger size fractions could be a result of their association with cellular or giant virus hosts. As an example of this phenomenon, the Mavirus virophage genome integrates into the genome of its cellular host *C. roenbergensis*, benefiting both virophage and the cellular host populations (74). As another example of intimate physical associations of virophages and their hosts, Sputnik virophage particles have been found embedded in *Mamavirus* fibrils which are a virion surface feature may play a role in entry into the giant virus host (75). Whether these examples are widely applicable to virophage-virus-host systems remains to be assessed, yet the high relative abundance of virophages in all > 0.45 μm size fractions is indicative of their potential influence on the *Mimiviridae* and their eukaryotic host communities.

We observed highly diverse and different virus communities in the > 0.45 μm size fractions, whereas the < 0.45 μm size fractions were similar and comprised primarily of Caudovirales. A recent study of virus attachment to particles suggested that up to 48% of viruses can adsorb to abiotic surfaces (76), with relevance to aquatic environments with high turbidity. Hamilton Harbour is a particle-rich environment with Secchi depths between 1.0-1.6 m at the 3 sites we sampled, and therefore virus attachment to suspended particles could be a significant factor in this environment. This could be one explanation for the higher relative abundances of viruses with < 0.45 μm virion sizes in the > 0.45 μm size fraction. However, virus attachment to particles was not directly assessed in this study. Overall, metagenomic sequencing of the smaller size fractions captured highly diverse Caudovirales communities, but the other virus families were absent or detected at only low relative abundances. Of Caudovirales for which hosts could be inferred, higher relative abundances of cyanophages were observed in the larger size fractions, potentially indicating replication within cells during ongoing infections. As noted previously, larger viruses including *Mimiviridae, Phycodnaviridae, Iridoviridae*, and *Poxviridae* were also captured in the > 0.45 μm size fractions. While some of these larger viruses were small enough to pass through the 0.45 μm pore-size filters, they may have been attached to cells, including non-host cells, or particles.

Our study demonstrates that analysis of virus communities using only the larger, > 0.45 μm diameter, size fraction can lead to underestimates of Caudovirales diversity, whereas analyzing only the smaller size fraction can lead to erroneously low estimates of diversity for many of the larger viruses such as *Mimiviridae, Phycodnaviridae, Iridoviridae*, and *Poxviridae*. Similarly, our data shows that examining only the smaller size fraction can lead to underestimation of virophage and cyanophage relative abundances that could, in turn, lead to researchers assuming their limited ecological importance in some environments. Furthermore, in this study, the smaller size fractions revealed relatively static virus community profiles, yet the community composition observed in the larger size fractions was much more variable. Clearly, virus community profiles were vastly different when comparing the free virus versus the cell- and particle-associated virus fractions, indicating that analysis of either size fraction alone will provide only a partial perspective of environmental viruses. Conversely, viruses detected in both size fractions may be potential candidates for further exploration to discern active viral infections in the environment. Given the considerable differences we observed in this study, with unique ecological patterns revealed for each size fraction, we recommend cautious interpretations of environmental virus community assemblages and dynamics when based on metagenomic data derived from different size fractions.

## MATERIALS AND METHODS

### Sampling sites and collection

Water samples were collected from three sites in Hamilton Harbour on September 20^th^, 2019. Site 1 (43°17.253” N, 79°50.583” W) is located in the middle of the harbour at its deepest point, approximately at the location of the mid-harbour site from our 2015 sampling (15). Site 3 (43°16.904” N, 79°52.527” W) is closer to the shoreline and corresponds to the nearshore site from our sampling in 2015, and Site 2 (43°17.135” N, 79°51.419” W) is located at the approximate mid-point between Sites 1 and 3. Surface water samples were collected in 20 L carboys using a bucket and were pre-filtered though 100 μm pore-size Nitex mesh to remove larger debris. A RBRmaestro^3^ multi-channel logger (RBR Ltd., Canada) was used to measure environmental physiochemical parameters including temperature, conductivity, dissolved O_2_, chlorophyll a, photosynthetically active radiation (PAR), salinity, and pH, and the Secchi depth at each site was also recorded.

Upon collection, water samples were covered and kept in the dark for immediate transportation and processing. After transportation to the lab, approximately 11 L of each sample was pressure filtered (100 mm Hg maximum pressure) through 0.5 μm nominal pore-size GC50 glass fiber filters placed atop 0.45 μm pore-size polyvinylidene fluoride (PVDF) filters. Following filtration, filters were stored at −20°C until DNA extraction. Filtrates, or water passing through the GC50 and PVDF filters, were treated with FeCl_3_ to precipitate viruses following the protocol of S. G. John et al. (43), except 0.45 μm pore-size PVDF filters instead of 0.8 μm pore-size polycarbonate (PC) filters were used to capture iron flocs after incubation. Filters with iron flocs were stored in the dark at 4°C until resuspension of viruses was performed according to the standard protocol (43). Resuspended viruses were transferred to polypropylene centrifuge tubes on top of a 1 mL sucrose cushion and pelleted via centrifugation at 182,000 x g for 3 hours at 4°C. Supernatants were decanted and 100 μl of TE buffer was added to each pellet which was covered with Parafilm to soften at 4°C before resuspension and nucleic acid extraction.

### DNA Extraction and Sequencing

For the > 0.45 μm size fractions, a quarter of each PVDF and GC50 filters was cut into pieces and transferred to separate screw-cap centrifuge tubes. Community DNA was extracted separately from each filter using the DNeasy Plant Mini Kit (Qiagen, Germany) following the manufacturer’s protocol, except that bead-beating instead of the TissueRuptor was used to disrupt samples. Approximately 0.25 g of each small (212 – 300 μm) and large (425 – 600 μm) glass beads were added to the centrifuge tubes with AP1 buffer and RNase A. The tubes were shaken in a FastPrep 96^®^ bead beater (MP biomedicals, USA) for 3 cycles of 90 seconds followed by 1 minute on ice between cycles. Final DNA concentrations were measured using a Qubit dsDNA HS Assay Kit (Life Technologies, USA). For each site, extracted DNA from the two filter types was combined in equal proportions based on DNA concentration to give a total volume of 40 μl and was stored at −20°C until they were sent for sequencing.

For the < 0.45 μm size fractions, virus pellets were resuspended by vortexing and community DNA was extracted using the QIAamp Viral RNA Mini Kit (Qiagen, Germany) following the manufacturer’s protocol. Final DNA concentrations were measured using a Qubit dsDNA HS Assay Kit (Life Technologies, USA), and DNA samples were sent for sequencing at the Microbial Genome Sequencing Centre (MiGS, Pittsburgh, Pennsylvania). Library preparation was performed using a Nextera Flex DNA Sample Preparation Kit (Illumina, USA) and sequenced from paired ends on the NextSeq 550 (2 x 150 bp) system (Illumina, USA).

### Metagenome Data Processing

Demultiplexing and adapter removal were performed by MiGS using bcl2fastq. Raw reads were processed according to C. N. Palermo et al. (15) with minor alterations to account for the different read lengths. Briefly, quality control was performed using Sickle version 1.33 (44) using a quality cut-off value of 30 and read length cut-off value of 50. Reads were assembled using IDBA-UD version 1.1.3 (45) with k values from 20 to 100 in increments of 20 and a minimum k-mer count of 2. BLASTx searches against the December 2019 NCBI-nr database (downloaded from ftp://ftp.ncbi.nih.gov/blast/db/FASTA/nr.gz) were performed using DIAMOND version 0.9.29 (46) with frameshift alignment and very sensitive modes activated. MEGAN6-LR version 6.14.2 (47) was used to annotate contigs using the November 2018 protein accession mapping file (downloaded from http://ab.inf.uni-tuebingen.de/data/software/megan6/download/welcome.html) with long read mode activated and bit score and e-value cut-offs of 100 and 10^-6^, respectively. Bowtie 2 version 2.3.5.1 (48) was used to map quality-controlled contigs back to the assembled contigs, and SAMtools version 1.9 (49) was used to extract the mapping formation for each contig. Table 1 summarizes the number of reads and contigs at each step in the processing pipeline for each sample.

### Metagenome Data Analysis

Data analysis methods followed those previously published in C. N. Palermo et al. (15). Briefly, contig relative abundances were estimated after normalizing by contig length and sequencing depth per sample. All virus contig assignments were categorized as one of the following based on NCBI taxonomic classifications: Caudovirales, virophages (*Lavidaviridae*), *Mimiviridae, Phycodnaviridae, Iridoviridae, Poxviridae*, other bacterial viruses, other dsDNA viruses, ssDNA viruses, and unclassified viruses. Taxonomic assignments for several contigs were less specific than information provided in the literature or NCBI’s taxonomy browser, and therefore these contigs were manually re-assigned to more specific groups (Table S1). Additionally, there were some contigs originally annotated as *Phycodnaviridae* that are now considered to be more closely related to *Mimiviridae* and are placed within the proposed subfamily “Mesomimivirinae” (32, 50). These contigs were manually re-assigned to the *Mimiviridae* category. *Marseilleviridae* and *Tectiviridae* were present at low relative abundances (< 0.01% and < 0.02%, respectively), and were therefore grouped together with the “Other dsDNA viruses”.

To assess the potential impact of Caudovirales on the prokaryotic community, Caudovirales contigs with inferred hosts that could be inferred from database information were extracted and analyzed separately. Host inferences were determined based on the name of the annotated virus. For example, a virus named *“Synechococcus* phage” was presumed to infect *Synechococcus*, whereas a virus named “Mediterranean phage” was presumed to infect an unknown host and was not included in the analysis of Caudovirales with inferred hosts. Caudovirales with inferred hosts were further classified as cyanophages or other bacteriophages.

Various packages in RStudio were used for data analysis and visualization as follows: virus relative abundances were presented as bar charts and bubble plots using the “ggplot2” package (51), unique contig annotations were compared between the size fractions using Venn diagrams generated using the “VennDiagram” package (52), Bray-Curtis dissimilarity analyses with the unweighted pair group with arithmetic mean (UPGMA) clustering were generated using the “vegan” and “dendextend” packages (53, 54), and finally, constrained correspondence analyses (CCA) were conducted using the “vegan” and “dendextend” packages (53).

### Data availability

DNA sequences generated in this study were deposited in the NCBI Sequence Read Archive (https://www.ncbi.nlm.nih.gov/sra) under the BioProject accession number PRJNA670357.

## ACKNOWLEDGEMENTS

DNA extraction and metagenomic sequencing was supported by an NSERC Discovery Grant (#RGPIN-2016-06022) awarded to S.M.S. Our appreciation to Dr. Bailey McMeans for the use of her sampling boat and RBRmaestro3 multi-channel logger. We thank Rose Mastin-Wood for assistance with sampling and sample processing.

## AUTHOR CONTRIBUTIONS

Conceptualization, C.N.P., D.W.S., and S.M.S.; methodology, C.N.P., D.W.S., and S.M.S; investigation, C.N.P., D.W.S., and S.M.S.; data curation, C.N.P. and S.M.S.; formal analysis, C.N.P. and S.M.S; validation, C.N.P. and S.M.S.; visualization, C.N.P. and S.M.S.; writing—original draft preparation, C.N.P. and S.M.S; writing—review and editing, C.N.P., D.W.S., and S.M.S.; supervision, S.M.S.; project administration, S.M.S.; resources, S.M.S.; funding acquisition, S.M.S.

